# Optogenetic operated probiotics to regulate host metabolism by mimicking enteroendocrine

**DOI:** 10.1101/2021.11.30.470589

**Authors:** Xinyu Zhang, Ning Ma, Wei Ling, Gaoju Pang, Tao Sun, Jing Liu, Huizhuo Pan, Meihui Cui, Chunli Han, Chun Yang, Jin Chang, Xian Huang, Hanjie Wang

**Affiliations:** School of Life Sciences, Tianjin University, Tianjin Engineering Center of Micro-Nano Biomaterials and Detection-Treatment Technology, Tianjin Key Laboratory of Function and Application of Biological Macromolecular Structures, 300072 Tianjin, China; School of Precision Instrument and Optoelectronics Engineering, Tianjin University, 300072 Tianjin, China; Laboratory of Synthetic Microbiology, Center for Biosafety Research and Strategy, School of Chemical Engineering & Technology, Tianjin University, 300072 Tianjin, China

## Abstract

The enteroendocrine system plays an important role in metabolism. The gut microbiome regulates enteroendocrine in an extensive way, arousing attention in biomedicine. However, conventional strategies of enteroendocrine regulation via gut microbiome are usually non-specific or imprecise. Here, an optogenetic operated probiotics system was developed combining synthetic biology and flexible electronics to achieve *in situ* controllable secretion to mimic enteroendocrine. Firstly, optogenetic engineered *Lactococcus lactis* (*L. lactis*) were administrated in the intestinal tract. A wearable optogenetic device was designed to control optical signals remotely. Then, *L. lactis* could secrete enteroendocrine hormone according to optical signals. As an example, optogenetic *L. lactis* could secrete glucagon-like peptide-1(GLP-1) under the control of the wearable optogenetic device. To improve the half-life of GLP-1 *in vivo*, the Fc domain from immunoglobulin was fused. Treated with this strategy, blood glucose, weight and other features were relatively well controlled in rats and mice models. Furthermore, up-conversion microcapsules were introduced to increase the excitation wavelength of the optogenetic system for better penetrability. This strategy has biomedical potential in metabolic diseases therapy by mimicking enteroendocrine.

## Introduction

The enteroendocrine cells (EECs) are scattered throughout the epithelium of the stomach, small and large intestines, collectively forming the largest endocrine organ^1^. EECs secretes about 20 gut hormones^2^ and plays an important role in the regulation of metabolic activities. EECs respond to a range of signals in the gut, including mechanical movements, nutrients, and microbial metabolites, which continuously regulate the secretion of gut hormones^1, 3^. The gut hormones have key roles in regulating food digestion and absorption, insulin secretion and appetite^4^. Metabolites of gut microbiome such as short-chain fatty acids can activate a number of cell types to regulatory hormone secretion^5, 6^. In recent years, there are some probiotics transplantation therapies to regulate and improve the enteroendocrine system, including microflora from healthy individuals^7, 8^ and genetically engineered bacteria^9^. Probiotics transplantation strategy has the advantages of convenient application, reducing the first pass effect of the digestive tract, and not requiring daily administration^10^. However, conversional probiotics transplantation strategies are usually non-specific or imprecise in the regulation of host metabolism.

Synthetic biology provides technical basis for the precise regulation of engineered bacteria. Various promoters were constructed responding to chemical or physical factors such as Isopropyl β-d-1-thiogalactopyranoside (IPTG)^11^, nisin^12^, NO^13^ and temperature^14^. Although these designs have good regulatory ability in specific application background, the ability is still greatly restricted^15^. Because of the complex physiological changes, seldom diseases have proper regulatory sensors, which limited the flexibility and generality of this strategy. In contrast, optogenetic tools can rapidly control gene expression with accurate spatiotemporal optical signal^16, 17^. Moreover, the light signal has minimal crosstalk with the human body. Therefore, to construct an optogenetic operated probiotics is suitable to secrete specific products accurately *in vivo*. Through the biosynthesis, the optogenetic operated probiotics is expected to controllably mimic endocrine in the intestine.

Here, we describe an optogenetic operated probiotics system (OOPS) for metabolism regulation by mimicking enteroendocrine. The system combines synthetic biology and flexible electronics to yield three major components that include optogenetically engineered probiotics and a wearable optical device. In this system, probiotics *Lactococcus lactis NZ9000* (*L. lactis*) was engineered with optogenetic promotor pDawn^18^ to secrete enteroendocrine hormone under the control of blue light. The wearable optical device could conveniently provide blue light for remote control of engineered *L. lactis*. In addition, considering further application of this system, better tissue penetration is demanded. Near-infrared (NIR) light is known to have greater tissue penetration than blue light^19^. For this reason, up-conversion microcapsules (UCMCs) were introduced in this system^20^. With the help of UCMCs, the engineered *L. lactis* could be operated by NIR light.

Glucagon-like peptide-1 (GLP-1) is a representative gut hormone released mainly from enteroendocrine L cells^21^. GLP-1 promotes the secretion of insulin in a glucose-concentration dependent manner, inhibits islet β cell apoptosis and help in weight control^22^. Therefore, GLP-1 has become a new target for blood glucose regulation in type 2 diabetes (T2D). The design and function of the system are shown in **Figure 1**. Furthermore, GLP-1 has a short half-life within the human body due to the degradation by enzyme DPP-IV^23^, limiting the suitability to deliver GLP-1 through systemic administration in clinical treatments. To enhance the stability, GLP-1 was C-terminally fused to IgG-Fc^24^ as more stable human GLP-1 (referred to as shGLP-1). We demonstrated the optogenetic operated probiotics system for GLP-1 and shGLP-1 secretion and glucose metabolism regulation effects in rat model of T2D. The strategy showed potential for convenient and precise metabolism regulation.

**Figure 1.**
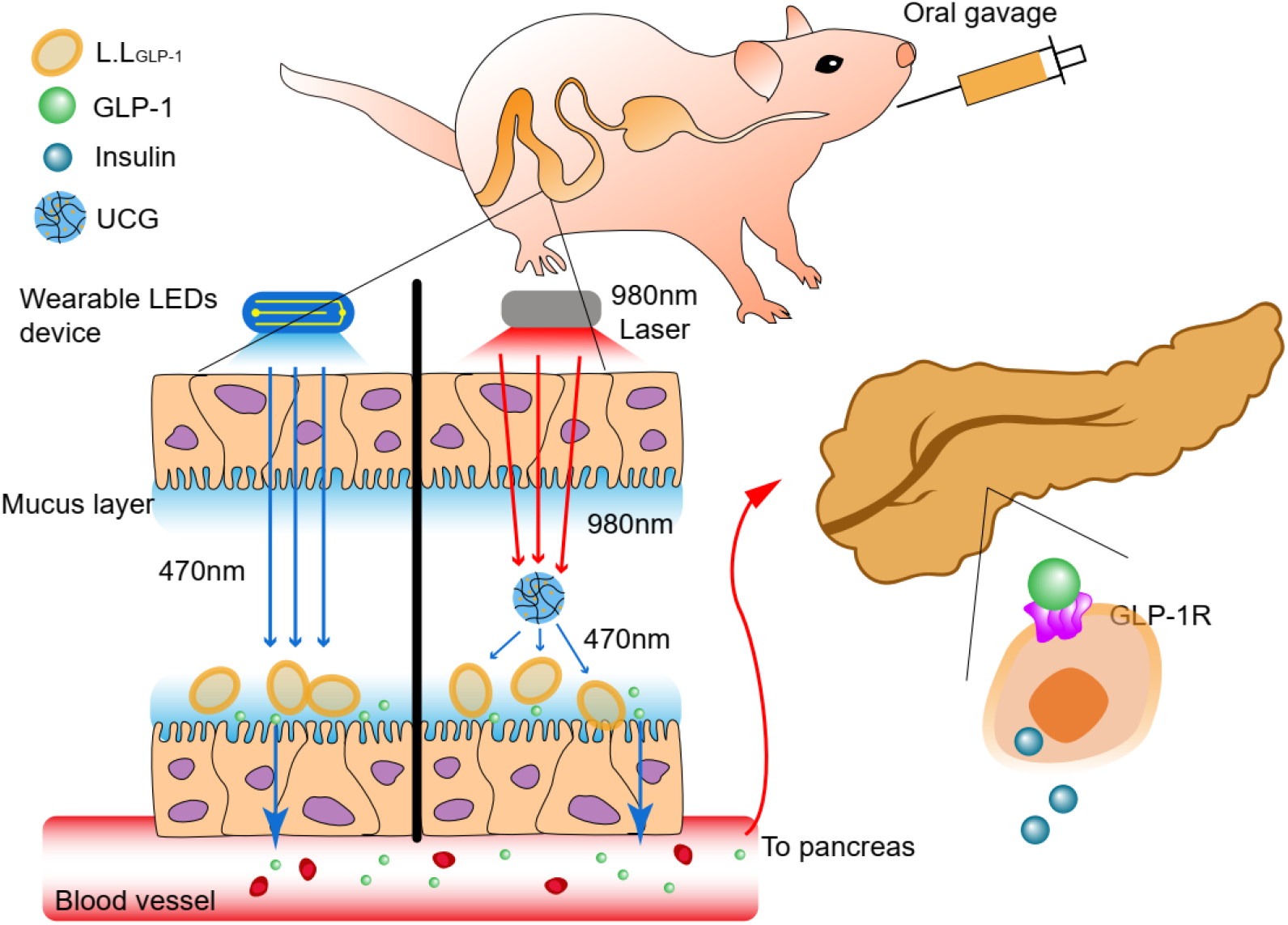
A schematic showing mechanism of glucose metabolism regulation via optogenetic operated probiotics system.

## Results

### Optogenetic probiotics function in the gut

The core component of the OOPS is optogenetic probiotics worked in the intestine. In order to demonstrate that pDawn can function in *L. lactis*, strain *L.L_GFP_* was constructed by transfecting plasmid pDawn controlling green fluorescent protein (GFP) expression **(Fig. 2a)**. After 30 mins of blue light stimulation of the strains in the logarithmic phase (OD=0.6), the fluorescence intensity of GFP increased progressively over time in the first 20 min **(Fig. 2b and 2c)**, proving the compatibility of pDawn in *L. lactis*. The pH of the medium decreased as the *L.L_GFP_* grow, leading to the fluorescence decay in 20-30min.

**Figure 2.**
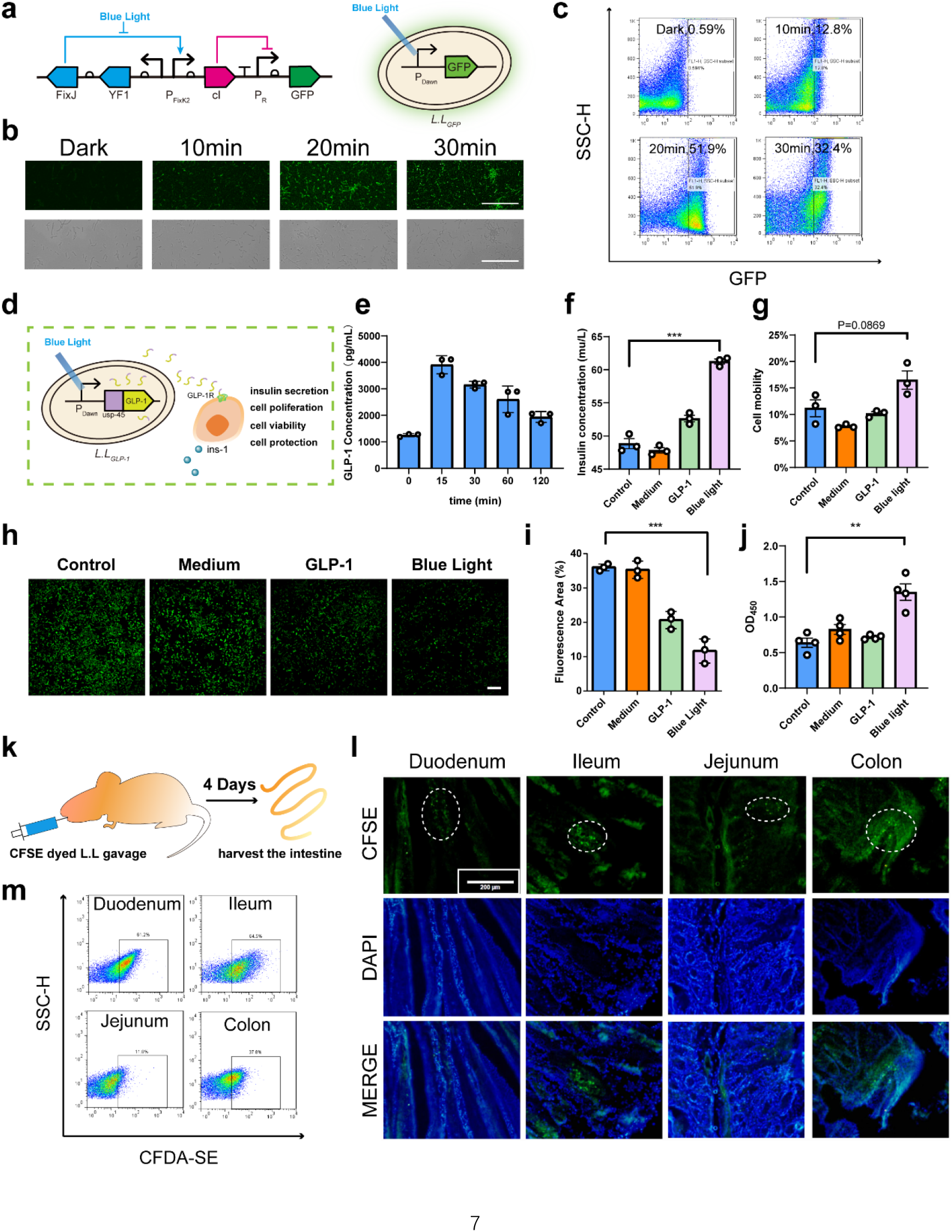
Construction and intestinal colonization of optogenetic probiotics. **a**. A schematic of *L.L_GFP_*. Plasmids were constructed and transformed to *L.lactis* (NZ9000) competent cells by electro-electroporation. **b**. Confocal microscope fluorescence images and flow cytometric analysis of blue light-induced GFP expression in *L.L_GFP_*. Scale bar: 50 μm. **c**. Flow cytometric analysis of blue light-induced GFP expression in *L.L_GFP_*. **d**. A schematic of *L.L_GLP-1_* secreting GLP-1 and triggering biological effects of ins-1 cells. **e**. GLP-1 secretion induced by blue light throughout 2 hrs measured by ELISA. **f**. Insulin secretion induced by GLP-1 in ins-1 cells. **g**. Cell mobility analysis of cell scratch experiment. **h. and i**. Oxidative stress experiment and its analysis via ImageJ. **j**. CCK8 cell proliferation assay of GLP-1 promotes the proliferation of ins-1 cells. (In **f-j**, cells in different group was added with same amount of PBS, *L.lactis* culture medium, 1ng/mL GLP-1 solution and *L.L_GLP-1_* after 1 h blue light induction, relatively.) **k**. Rats received 4 days of CFSE dye-stained *L.L_GLP-1_* gavage for intestinal engraftment prior to harvest of the intestinal tract on the fifth day. **l**. Fluorescence microscope images of CFSE dyed *L.L_GLP-1_* in different intestinal segments. Scale bar: 200 μm. **m**. Flow cytometric analysis of CFSE dyed *L.L_GLP-1_* from different intestinal segments. All data are mean ± SEM. P values were calculated by Student’s t-test (***p < 0.001, **p < 0.01, *p < 0.05 criteria).

Subsequently, we synthesized the gene sequence of usp45 (a lactobacillus secrete signal peptide^25^) fused GLP-1(7-37) and constructed light controlled GLP-1 secreting *L. lactis* (*L.L_GLP-1_*, **Fig. 2d, Fig. S2a)**. The growth curves of *L.L_GLP-1_* **(Fig. S2a)** under different light/temperature conditions were measured. *L.L_GLP-1_* of blue light/dark group in 30°C reached the maximum range of turbidity (the dotted line) after 4 hours of cultivation, while under dark conditions at 37°C (to stimulate body temperature) the *L.L_GLP-1_* clone entered the stationary phase, indicating blue light wouldn’t interfere with the growth of *L.L_GLP-1_* but the temperature would inhibit its growth. Western blot experiments showed that GLP-1 was successfully synthesized in the bacteria after light induction for 1 h **(Fig. S2b)**. ELISA also confirmed GLP-1 secretion in the supernatant **(Fig. 2e)**. Affected by the burden synthetic circuits and instability^26, 27^, we found concentration of GLP-1 peaked during the initial induction phase and decreased gradually. To demonstrate that GLP-1 secreted by LL was biologically active and available to mimic enteroendocrine, we used ins-1, a rat islet tumor cell line, to validate a range of biological effects^22, 28^. For promoting insulin secretion, we co-cultured *L.L_GLP-1_* with ins-1, a rat islet tumor cell line, followed by blue light stimulation (30 mW/mm^2^) for 1 h. Insulin secreted by ins-1 into the cell medium detected by ELISA showed a significant improvement **(Fig. 2f)**. Under cell scratch experiments, GLP-1 showed slight enhancement in cell migration ability **(Fig. 2g and Fig. S3)**. Subsequently, the medium supernatant of *L.L_GLP-1_* induced by blue light for 15 minutes was added to ins-1 to verify its role in protecting islet β cell from oxidative stress induced by hydrogen peroxide **(Fig. 2h and 2i)**. In addition, significant cell proliferation induced by GLP-1 secretion under optogenetic control was observed compared to control using CCK8 (Cell Counting Kit-8) detection **(Fig. 2j)**.

Following the above in-vitro measurements, the exist and retention of engineered *L. lactis* was detected. Carboxyfluorescein succinimidyl amino ester (CFSE) dyed *L.L_GLP-1_* was delivered to the intestinal tract of rats by oral gavage **(Fig. 2k)**. Flow cytometric analysis and intestinal tissue fluorescence microscope images showed its retention in duodenum, jejunum, ileum, and colon **(Fig. 2l and 2m)**. Moreover, *L.L_GLP-1_* persisted in the intestinal tract for 7 days after gavage **(Fig. S2c)**. The successful construction and long-time retention of *L.L_GLP-1_* shows the potential of the optogenetic probiotics for regulating glucose metabolism *in vivo*.

### Construction of wearable optical device

A wireless-controlled, flexible optical device with a weight of only 4.798 g was fabricated to trigger the *L.L_GLP-1_* using blue light **(Fig. 3a, Fig. S4 and Supplementary Videos)**. A Bluetooth module was used to implement wireless control through a smart-phone application. Based on the physiological structure of the rat intestine, four LED light sources have been arranged squarely to cover the entire length of the small intestine. Each source consisted of four mini-LEDs to ensure the necessary light intensity **(Fig. 3b and 3c)**. The optical and electrical properties of the device were also characterized. Mini-LEDs with a size of 0.8 × 0.3 mm^2^ showed a forward current of 9 mA under a voltage of 3.3 V **(Fig. 3d)**. The emission wavelength of the device has been measured to be 462 nm, which complies with the activation wavelength of promoter pDawn **(Fig. 3e)**. In order to avoid the high temperature caused by continuous stimulation, graphene membranes were modified to the backside of the flexible printed circuit board to facilitate heat dissipation. Also, the device was allowed to work in a pulse-mode with a peak working current of ~60 mA to regulate the temperature of the LED chip within an acceptable range less than 40 °C **(Fig. 3f and 3g)**. Tissue penetration of LED photons was also simulated according to optical absorbance and scattering parameters of the 470 nm wavelength in different tissues using a Monte Carlo simulation software described in details in the supporting information. The simulation result showed that the intensity in the gut, after penetrating the rat external tissues, was around 1.5 mW/mm^2^ **(Fig. 3h and 3i)**, which is greater than the minimum excitation intensity required to activate promoter pDawn (1 nW/mm^2^)^18^.

**Figure 3.**
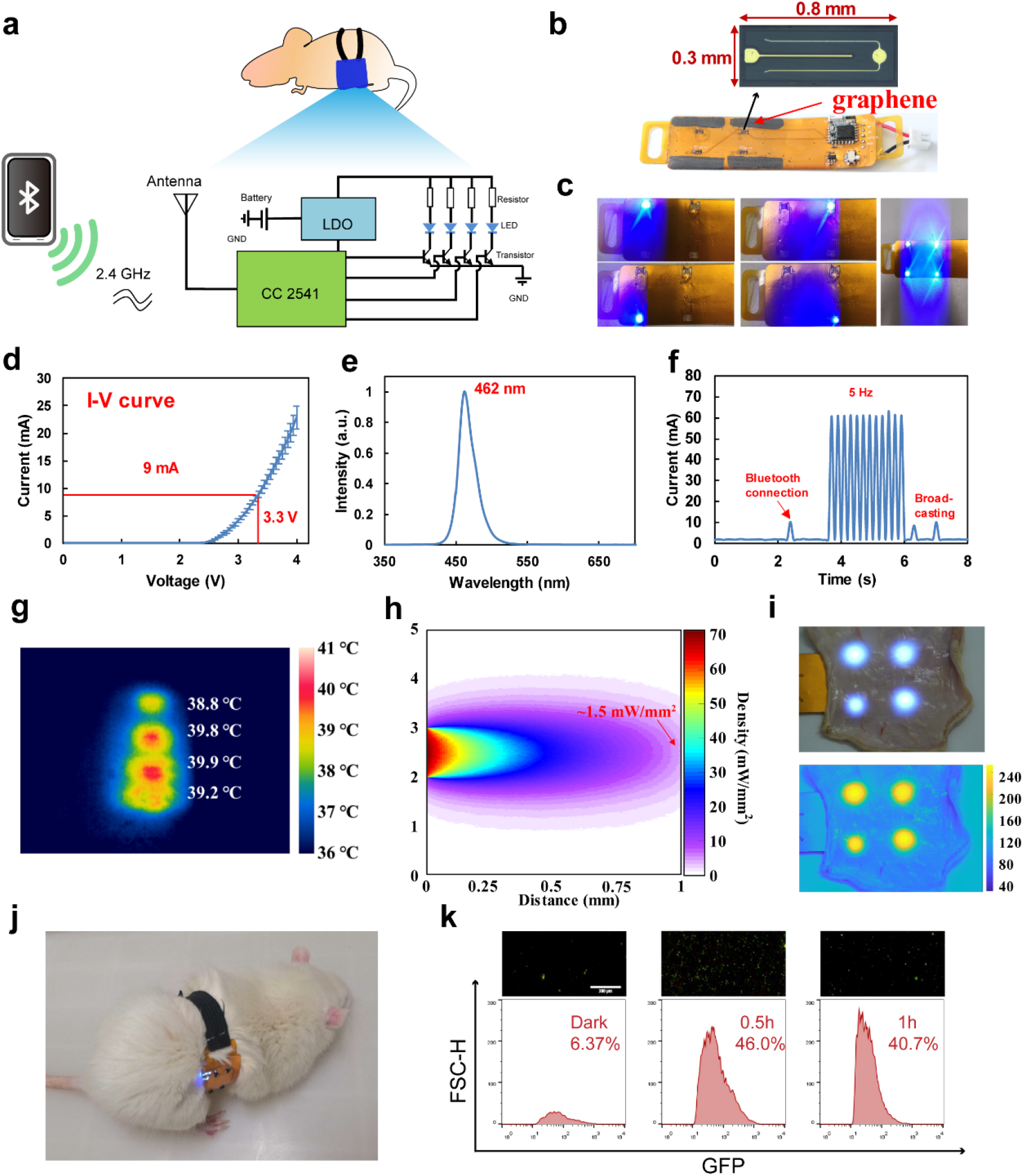
Characterization of wearable optical device. **a**. A schematic of the electrical operation principles of the device. The device was connected to a smart-phone via a Bluetooth module for wireless control of mini-LEDs. **b**. Enlarged images of a mini-LED and a flexible optical device. Graphene membranes were deposited onto the backside of the device to facilitate heat dissipation. **c**. Images of a wireless-controlled flexible optical device showing separate control of each LED. **d**. Voltage-current characteristics of mini-LEDs. *n* = 6, error bars indicate means ± SD. **e**. The electroluminescent spectra of the device with a peak emission wavelength of 462 nm. **f**. Power consumption of the device when working in a pulse mode. **g**. An infrared image of the device showing the temperature distribution of mini-LEDs. The peak temperature was 39.9°C. **h**. Simulation of light penetration depth in rat skin tissue using the device. **i**. A representative image and corresponding grayscale image of the blue light of the device after penetrating the skin tissue. **j**. A freely moving rat equipped with the device. **k**. Microscope fluorescence images and flow cytometric analysis of *L.L_GFP_* colonized in the intestinal tract induced by the device. Scale bar: 200μm.

The feasibility of the OOPS with wearable optical device for *in vivo* glucose regulation was demonstrated. Rats were given *L.L_GFP_* by oral gavage for 4 days, followed by shaving hairs on the belly before equipping the LED device to reduce the light loss **(Fig. 3j and Fig. S5)**. The intestines of the rats were harvested after 0 h, 0.5 h and 1 h of blue light stimulation. Intestinal contents and villi were scraped to collect bacteria for analysis. Microscope fluorescence micrographs and flow cytometric analysis showed that the LED device successfully induced GFP expression of *L.L_GFP_* in the intestinal tract **(Fig. 3k)**.

### Metabolism regulation via wearable optical device

Based on optogenetic *L. lactis* and optical device, the ability of OOPS to regulate glucose metabolism in vivo was evaluated in the T2D rat model **(Fig. 4a)**. 7-week-old rats were firstly fed with a high-fat diet (HFD) for 2 months to produce insulin resistance **(Fig. S6a)**. Each rat was then injected with Streptozotocin (STZ-1) and finally developed symptoms of T2D indicated by a high glucose levels of 20mmol/L on average compared with 5mM for normal rats and 7mM for only HFD rats **(Fig. S6b)**. Rats in Blue group (treatment group) then received *L.L_GLP-1_* gavage for 4 days. Rats were then equipped with wearable optical device and received blue light (30 mW/mm^2^, 5 Hz) for 1 h per day. Blood samples were collected right after stimulation, and blood insulin and GLP-1 concentrations were both almost twice than control **(Fig. 4c and 4d)**. Glucose was found to be reduced from 24.4 mmol/L to 20.9 mmol/L during stimulation, while the glucose levels increased slightly from 24.9mmol/L to 27.2mmol/L in average for the control **(Fig. 4e)**. Random blood glucose of treated rats showed gradually decrease from 27.4mmol/L in day one to 15.1mmol/L in day 6 **(Fig. 4f)**. After daily treatment for one week, the random blood glucose of blue group reduced by 7 mmol/mL as compared with the control group **(Fig. 4g)**. However, the intraperitoneal glucose tolerance test didn’t show significant difference in glucose tolerance **(Fig. 4h and 4i)**, which may due to insufficient GLP-1 secretion and treatment period. In addition to long-term hyperglycemia, the high-fat diet and STZ-1 induced characteristic hyperlipidemia in T2D rats **(Fig. 4j(1))**. Blood lipids remained normal in the blue light group, and began to rise after treatment was discontinued **(Fig. 4j and 4l)**, indicating that GLP-1 also plays an important role in lowering blood lipids^29^.

**Figure 4.**
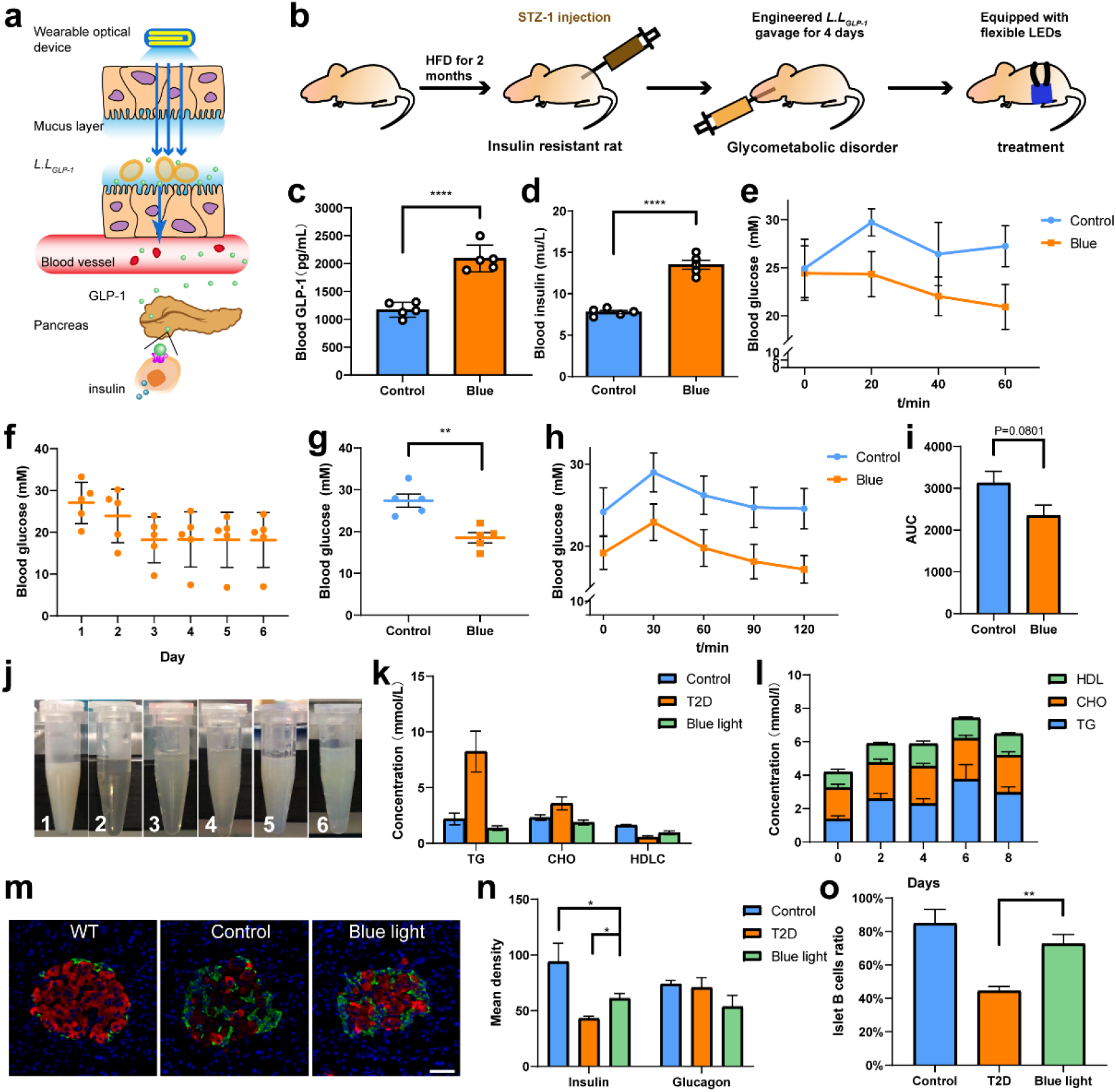
Glucose metabolism regulation via wearable optical device on T2D rats. **a**. A schematic of the working principles of glucose metabolism regulation using a wearable optical device. **b**. A schematic of the timeline of animal experiment. **c** and **d**. Blood GLP-1 and blood insulin variation between blue light group and control group (Detected by ELISA). **e**. Short-term time course of blood glucose change of rats in a blue light group during stimulation. **f**. Random blood glucose of rats in the blue light group over the course of 6 days, each rat received 1 h of blue light stimulation per day. **g**. Blood glucose in the blue light group and control group after 1 week. **h**. Intraperitoneal glucose tolerance test. **i**. Area under the curve (AUC) analysis of the intraperitoneal glucose tolerance test in **h**. **j**. Changes in serum at the end of treatment. The serum of rats in the control group is chylous (**1**). **2-6** Serum of rats in blue light group 0, 2, 4, 6, 8 days after stopping blue light stimulation. **k**. Blood lipid test of normal rats (WT), control and blue light group rats at the end of the treatment. **l**. Blood lipid test of blue light group rats at 0, 2, 4, 6, and 8 days after stopping blue light stimulation. **m**. Fluorescent images of islet (Blue: DAPI, Red: insulin of islet β cells, Green: glucagon of islet α cells. Scale bar: 40 μm). **n** and **o**. Fluorescence analysis of islet fluorescence micrographs. **n**. Mean fluorescence intensity represents the activity of glucagon in islet α cells and insulin in islet β cells. **o**. The proportion of islet cells in the islet. All the data are mean ± SEM. P values were calculated by Student’s t-test (***p < 0.001, **p < 0.01, *p < 0.05 criteria). Sample size: n= 7 (**c**), n=5 (**d-l**).

After all the above measurements were completed, rats were sacrificed for tissue section analyses. Islet fluorescence staining showed higher activity and proportion of islet cell β **(Fig. 4m-4o and Fig. S7)**. Hematoxylin and eosin (H&E, **Fig. S8**) stained slices of pancreas tissues show common vacuolization of the cytoplasm (black arrow), rare punctate necrosis and nuclear fragmentation (yellow arrow) of islet cells in the control group. That there is only a decrease of islet cell cytoplasm in the blue light group indicates the protective effect that GLP-1 exerts on islet cells. HE staining sections of other tissues showed no significant adverse effects **(Fig. S8)**. Thus, we conclude that the animal experiment demonstrates that the OOPS lowers blood glucose by 10%-20% and improves the blood lipid metabolism, suggesting the potential of this strategy for glucose metabolism regulation by mimicking GLP-1 enteroendocrine.

### Optimizing the stability of GLP-1

Due to the short half-life of GLP-1, mimicking GLP-1 enteroendocrine with optogenetic probiotics might be deficient in long-term effects, and thus limiting the biomedical translation prospects of this strategy. To optimize the stability of GLP-1, we constructed another *L. latics* secreting clone (*L.L_sh_*), that expressing more stable human GLP-1 (shGLP-1) under blue light control **(Fig. 5a, Fig. S9a)**. *L.L_sh_* exhibited continuous increase in shGLP-1 secretion even after 60 mins of blue light exposure. In contrast, GLP-1 secretion reached the maximum value after 20 mins of light exposure with *L.L_GLP-1_* **(Fig. 5b)**, indicating that introduction of *L.L_sh_* can effectively extend the working periods of the bacteria. To achieve more precise GLP-1 delivery control, we also explored the relationship between the amount of *L.L_sh_* gavage and GLP-1 secretion. When the amount of *L.L_sh_* gavage varied from 1 mL to 4 mL, the concentrations of shGLP-1 measured after light stimulation of 60 min were between 7.9 and 8.5 pM, suggesting that increased *L.L_sh_* would not significantly increase the secretion of shGLP-1**(Fig. 5c)**. To compare the effectiveness of using light stimulation of *L.L_sh_* to regulate insulin levels, referred to as *L.L_cons_*, another modified *L. latics* that can spontaneously and constantly secret shGLP-1 without optogenetic control has been adopted **(Fig. S9b)**.

**Figure 5.**
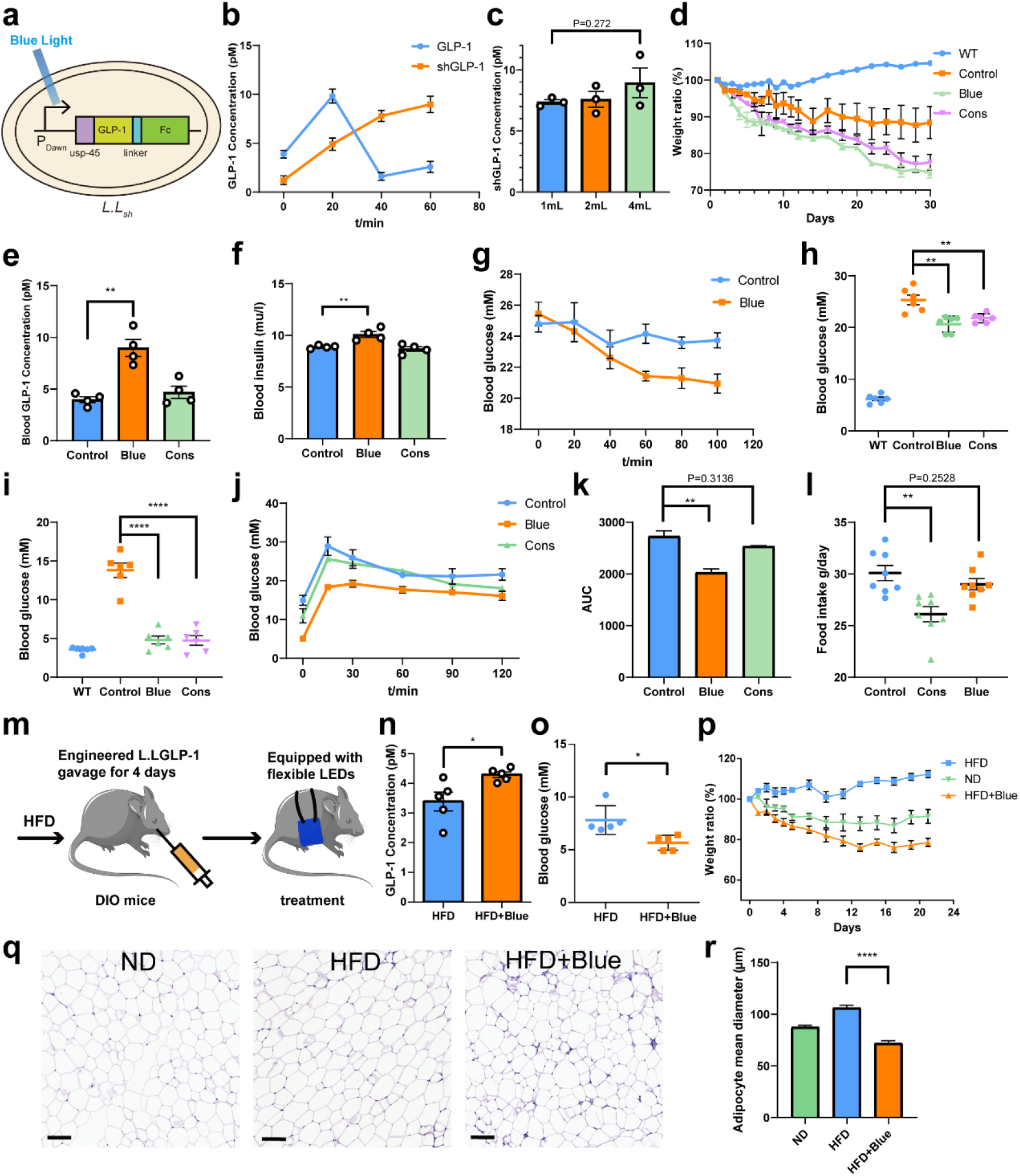
shGLP-1 enhances the stability of GLP-1. **a**. A schematic of the design of *L.L_sh_*. **b**. Time course of GLP-1 and shGLP-1 concentrations in vivo under blue light stimulation. **c**. Blood GLP-1 concentration under blue light stimulation after different doses of *L.L_sh_* gavage. **d**. Weight changes in rats during one month of treatment (WT, Control, Blue and Lac group corresponding to normal, diabetic, *L.L_sh_* and *L.L_lac_* gavage rats, respectively). **e**. and **f**. Blood GLP-1 and insulin concentrations of rats were tested right after blue light stimulation. **g**. Time course of blood glucose concentration of rats under blue light stimulation. Light was turned on right after blood glucose t0 was measured. **h** and **i**. Random blood glucose (**h**) and fasting blood glucose (**i**) concentration of rats in different groups. **j**. Intraperitoneal glucose tolerance test of rats analyzed on the 20^th^ day of treatment. **k**. Area under the curve (AUC) analysis of the intraperitoneal glucose tolerance test in (**j**). **l**. Food intake of rats in different groups. **m**. A schematic of the timeline of animal experiment on DIO mice. **n** and **o**. Blood GLP-1 and glucose concentration change during blue light stimulation in DIO mice. **p**. Weight changes in DIO mice (HFD, high fat diet and ND, normal diet). **q**. Representative hematoxylin and eosin (H&E)-stained pictures of adipose tissue of DIO mice. Scale bar: 100μm. **r**. Mean adipocyte diameter (μm). All the data are mean ± SEM. P values were calculated by Student’s t-test (***p < 0.001, **p < 0.01, *p < 0.05 criteria). Sample size: n=3 (**c**), n≥4 (**d-p**).

Diabetic rats received *L.L_sh_* or *L.L_cons_* gavage for 4 days, serum was collected on day 0, 2, 4, 6 and 8 after gavage, the levels of inflammatory factors (IL-2, IL-6, TNF-α) were measured by ELISA to confirm the immune safety of gavage **(Fig. S10)**. The *L.L_sh_* gavage group was treated for 30 days by receiving 1 h of blue light stimulation (30 mW/mm^2^, 5 Hz) per day. Moreover, shGLP-1 could act on the brain and intestines, inhibits intestinal peristalsis and gastric emptying **(Fig. S9c-f)**, food intake was inhibited in the treated rats, and thus cause weight loss, which is beneficial to the treatment of T2D **(Fig. 5d and 5l)**. During blue light stimulation, the serum GLP-1 concentration significantly increased, and insulin secretion was promoted to reduce blood glucose by 6.05±2.45 mmol/L **(Fig. 5e-5g)**. Compared with continuously secretion of shGLP-1, optogenetic control would reduce the burden of *L. latics*, thus higher concentration of shGLP-1 was detected in blue group. Random blood glucose test also showed reduced blood glucose in blue and cons group **(Fig. 5h)**. On the 20^th^ day of treatment, we performed an intraperitoneal glucose tolerance test. Blue light treatment markedly reduced fasting glucose level **(Fig. 5i)** and exhibit better glucose tolerance than *L.L_cons_* **(Fig. 5j and 5k)**. Because GLP-1 has multiple physiological functions, we also examined the cardiovascular effects of the treatment. The treatment successfully restored TG and TC concentration to normal level in the first days, no significant change was observed in heart rate and blood pressure **(Fig. S11)**. At last, to verify the effect of the treatment on the intestinal flora, a microbiota sequencing experiment was performed, we did not find any adverse effects after a month-long exposure to *L. lactis* on other intestinal flora **(Fig. S12)**. We interpret this as evidence of the biosafety of OOPS.

Subsequently, to evaluate the adaptability of other animal models, the system has been used on diet-induced obesity (DIO) mice. Each mouse was given 500μL *L.L_sh_* for 4 days followed by continuous blue light stimulation for 1 hour each day during the entire experimental period of 3 weeks **(Fig. 5m)**. Daily blue light stimulation exhibited increased blood GLP-1 concentration and lower blood glucose concentration compared to the untreated mice **(Fig. 5n and 5o)**, while the insulin level and glucose tolerance maintained **(Fig. S13)**, indicating other ways of blood glucose handling beside the insulinotropic effect of shGLP-1. However, the treated mice exhibited significant weight loss **(Fig. 5p)** after three weeks of treatment. The mice were sacrificed to acquire adipose tissue for H&E staining analysis **(Fig. 5q and 5r)**. The result indicated reduced adipocyte diameter due to the SIRT1-dependent lipolytic and oxidative capacity effect of shGLP-1^30^.

### Probiotics operation via up-conversion microcapsules

Continuous exposure to strong blue light might cause cytotoxic effect^31^. Besides, poor tissue penetration of blue light further limits its clinical application. To side-step these issues, up-conversion materials were used for blood glucose metabolism management under near-infrared control **(Fig. 6a and 6b)** ^32, 33^. Using up-conversion microrods as an energy transfer relay, near-infrared light (980 nm) with deeper penetration than blue light **(Fig. S14)** is converted to blue light (475 nm) in vivo **(Fig. 6c and 6d, Fig. S15)**. Then the blue photons stimulated gene expression of plasmid pDawn in *L.L_GLP-1_*. However, because delivering up-conversion microrods to the intestinal tract is cytotoxic and leads to excessive dispersion of the microrods that would reduce the intensity of the glow, the microrods were encapsulated in a biocompatible microcapsule^34^ **(Fig. S16)** using microfluidic technology **(Fig. 6e and 6f)**. The microdroplets showed high stability, low permeability at different pH over 20 hours **(Fig. 6g and 6i)**. Then ICG (Indocyanine Green) encapsulated microcapsules were delivered to the intestinal tracts of rats. These were used to monitor in vivo distribution of the microdroplets by imaging the intestinal tract and the feces every 4 hours using an in-vivo imaging system **(Fig. 6h)**. The microcapsules were localized in the small intestine in the first 4 hours and started to be expelled from the body at around 8 hours after gavage.

**Figure 6.**
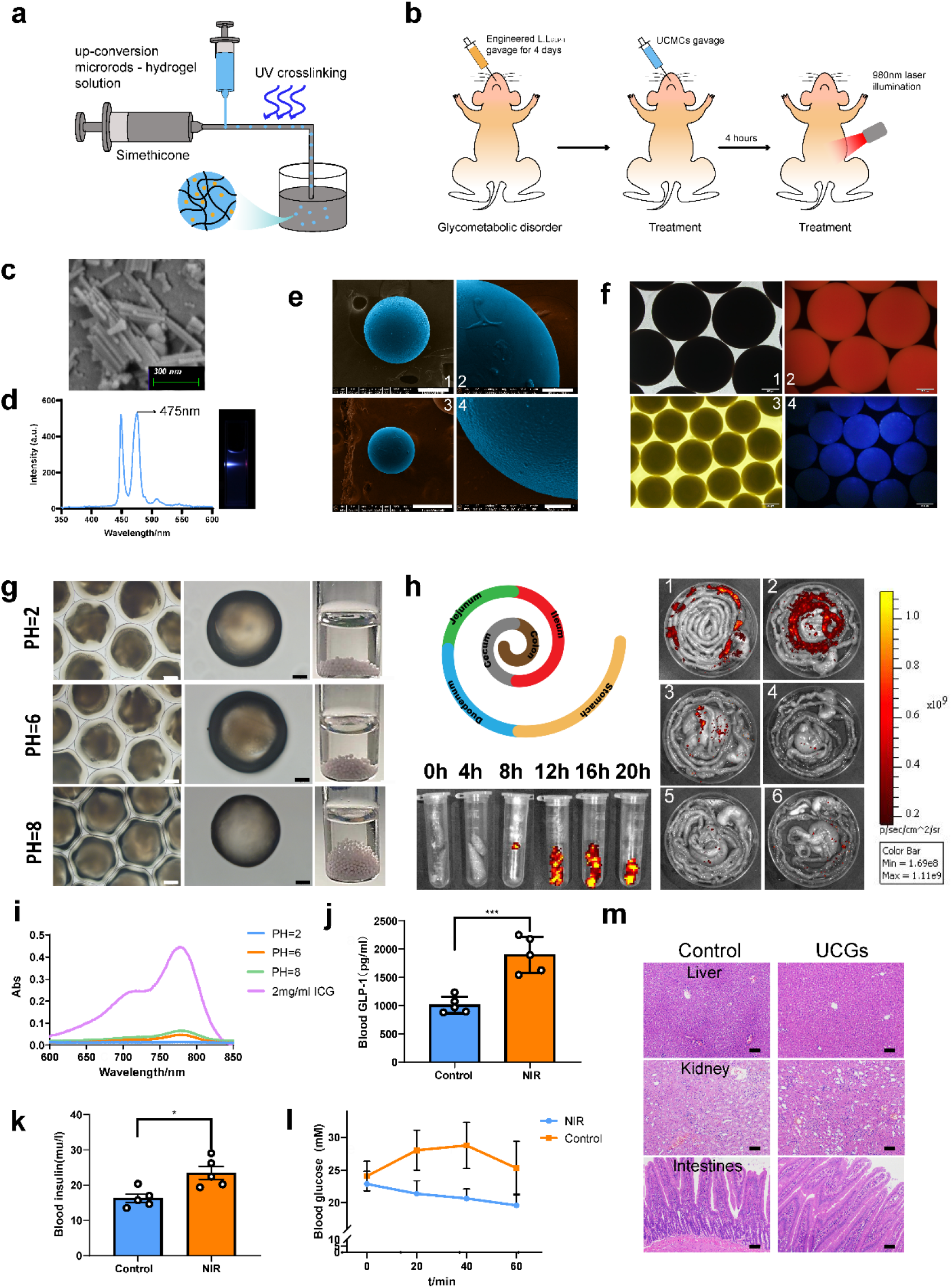
*L. Lactis* operation via up-conversion microcapsules. **a**. A schematic of the synthesis of up-conversion microrods encapsulated by micro-droplet gels to form UCMCs. **b**. Illustration of the use of microrods to deliver near-infrared light through the skin and its absorption in the gut. Upon absorption by the up-conversion microrods, NIR photons are converted to blue photons where they activate the optogenetic translation of GLP-1 in *L. lactis*. **c**. Scanning electron microscope (SEM) images of up-conversion microrods. Scale bar: 300 nm. **d**. Emission wavelength of up-conversion microrods. **e**. SEM images of empty microcapsules (**1** and **2**) and encapsulating up-conversion microrods (UCMCs, **3** and **4**). Scale bar: 200 μm (**1**), 50 μm (**2** and **4**), 300 μm (**3**). **f**. Microphotograph of UCMC encapsulating ICG excited by 780 nm wavelength light (**1** and **2**) and UCMCs excited by 980 nm near-infrared (NIR) light (**3** and **4**). Scale bar: 200 μm. **g**. Stability of UCMCs in different pH environments measured after 20 hrs exposure. Scale bar: 200 μm (white); 100 μm (black). **h**. Enterodynamic experiment performed by *in vivo* imaging using UCMCs encapsulating ICG placed in the gut by gavage. The UCMCs stay intact in the intestinal tract and is expelled from the body within 12 hrs after gavage. **1-6** Distribution of gel encapsulating ICG in the intestine within 0-20 h. **i**. UV absorbance curve of ICG in solutions of different pH. **j** and **k** Blood GLP-1 and blood insulin variation between NIR group and control group (detected by ELISA). Each rat received 24 mW/mm^2^, 5 Hz NIR stimulation for 1 h per day. **l**. Blood glucose regulation under NIR stimulation. **m**. Hematoxylin and eosin (H&E)-stained slices of liver, kidney and intestines of the control group and UCMCs gavage group (Scale bar: 100 μm). All the data are mean ± SEM. P values were calculated by Student’s t-test (***p < 0.001, **p < 0.01, *p < 0.05 criteria). Sample size: n≥ 5.

To validate the up-conversion control in vivo, *L.L_GFP_* and UCMCs were delivered to the rats by gavage. GFP was successfully expressed after 0.5 hour and 1 hour of exposure to NIR laser stimulation **(Fig. S17)**. Subsequently, the *L.L_GLP-1_* and the UCMCs were delivered to the diseased rats by gavage. One hour after stimulation under a near-infrared laser, more than 88.9% higher in blood GLP-1 and 44.3% higher in insulin concentrations were detected in the treated group through ELISA right after the stimulation **(Fig. 6j and 6k)**. The blood glucose of rats in NIR group reduced by 3.3mmol/L in comparison with almost unchanged blood glucose in the control group **(Fig. 6l)**. H&E-stained slices of liver, kidney and intestines of the treated rats revealed no conspicuous damage **(Fig. 6m)**, indicating that the up-conversion optogenetic nano system affords a biosafe regulation of glucose metabolism in vivo.

## Discussion

A system for metabolism regulation via optogenetic probiotics has been put forward, which introduced synthetic biology to design optogenetic *L. lactis* and adopted flexible electronics techniques and rare-earth nanotechnology to control *L. lactis*. Considering the flexibility of synthetic biology design and the convenience of optogenetic control, this strategy theoretically has universality in biomedical applications. Due to the intestinal function, the effects of engineered probiotics administration can mimic the enteroendocrine pathway. Therefore, the strategy provides a new intervention method for the metabolic regulation which the enteroendocrine system involved in, and has prospects for the treatment of metabolic diseases.

Despite a complete solution has been given here, it is worth to be mentioned that several challenges remain. Firstly, every segment of enteroendocrine and corresponding metabolism is complicated, and may involve the interaction of multiple organs, cells, and signal pathways^2, 35^. As a metabolism regulation strategy, the development of this strategy depends on the achievements of enteroendocrine mechanism research. At the same time, we expect that the OOPS can help in the mechanism research of enteroendocrine in the future. Otherwise, by secreting shGLP-1, the blood glucose of T2D rats decreased by 6mM, but still far from the normal level. This may be caused by resistance of the *L. lactis* to prolonged blue light stimulation, leading to limited concentration of blood shGLP-1. We hypothesize that this limitation can be optimized to some extent by introducing genetic circuits^36, 37^. Besides, *L. lactis* is a prokaryote, it can produce only a limited number of hormones, and the products tend to accumulate in the cytoplasm limited by the secretion capacity. The follow-up work needs to focus on synthetic biology to enrich product types^38^ to better mimic enteroendocrine.

In brief, we have proposed a new strategy that accurately and effectively regulates metabolism by mimicking enteroendocrine. This strategy is based on optogenetic probiotics and involves two technologies to assist in light control: wearable devices and up-conversion microcapsules. This strategy is precise and controllable for metabolic intervention, and has potential universality. It might bring new opportunities for the treatment of metabolic diseases, and is expected to inspire the research on the relationship between gut microbes and the host. For this goal, in the future, we will conduct more in-depth studies on the interaction between the engineered probiotics and the host.

## Materials and Methods

### Construction of strains

Plasmid pDawn was constructed according to the method of Ohlendorf et. al. Gene sequence of signal peptide usp45, rat GLP-1 and Fc of IgG in rat was found in NCBI. The recombinant plasmid was constructed by PCR using AceTaq® DNA Polymerase (),and finally transformed to *L. lactis NZ9000* competent cells by electro-electroporation.

### Flow cytometry analysis

In-vitro induced bacteria were centrifuged at 8000 rpm for 5 mins and washed twice with PBS. Rats were sacrificed, and the intestine was removed and segmented. The intestinal mucosa was collected and resuspended in PBS at a volume ratio of 1:1. The precipitate was removed by centrifugation at 2000 rpm for 5 mins, and the bacteria were collected by centrifugation at 8000 rpm for 5 mins. Both sediments were fixed with 1% paraformaldehyde for 20 mins followed by PBS washing. The sediment was then resuspended in 1 mL PBS before flow cytometry analysis. Flow cytometric analysis was performed using a FACSCalibur instrument (Becton Dickinson, Sunnyvale, CA). Data were analyzed on Flowjo VX.

### Cell experiment

Cellular cultures of ins-1 were incubated in RPMI-1640 medium in 12-well cell culture plates to a proper density. *L.L_GLP-1_* in the logarithmic phase was added in 100 μl aliquots to ins-1 per well. Then, 100 μl bacterial medium and 1 ng/mL GLP-1 solution were added to other wells as negative control groups. Co-culture lasted for 1 h under 25 mW/cm2 blue light stimulation. The supernatant was then collected and tested using an insulin ELISA kit. For cell proliferation assay, ins-1 was incubated in a 96-well cell culture plate at a density of 2000 cells per well. *L.L_GLP-1_* in the logarithmic phase was induced under 25 mW/cm^2^ blue light for 1 h. The supernatant was collected and filtered with a 0.2 μm filter membrane, then added as cell-free supernatant to ins-1 at 10 μl per well. Next, 10 μl bacterial medium and 1 ng/ml GLP-1 solution were added to other wells as negative control groups. The culture plates were incubated in a cell incubator for 12 h. A CCK-8 cell viability kit was used to monitor cell growth by measuring the absorbance at OD450 using a microplate reader. Oxidative stress protection was detected by ROS-ID® Hypoxia/Oxidative stress detection kit(ENZ-51042-0125, Enzo Life Sciences Inc., Switzerland). The supernatant of *L.L_GLP-1_* after 1 hour blue light induction and incubated for 24 h in 12-well plates, followed by ROS induction, confocal and flow cytometry detection as recommended in the manual.

### Validation of intestinal *L.L_GLP-1_*

*L.L_GLP-1_* was co-cultured for 30 mins with 50 μM CFSE at a volume ratio of 10:1, then centrifuged for 3 mins at a speed of 8000 rpm. Unbound dye was removed by PBS wash for 2 cycles. The CFSE dyed *L.L_GLP-1_* was delivered to rats by continuous oral gavage for 4 days. On the fifth day, rats were sacrificed and harvested the intestine. Samples were soaked in paraformaldehyde for 1 day and divided into segments identified as duodenum, jejunum, ileum (into two parts), and colon. Each segment was divided into two sections: one section was immobilized in the embedding fluid for use in frozen sections, the other was dissected and the mucosa scraped to harvest cells for flow cytometric analysis.

### Construction of the wearable optical device

Low footprint, packaged off-the-shelf components were chosen as the basis of the circuit. High-brightness mini-LEDs (S-32CBMUD, Sanan Optoelectronics) were mounted to a flexible printed circuit board using low-temperature Ag solder paste. A low-power microcontroller (CC2541, Texas Instruments) with Bluetooth capability controlled the working sequence of LEDs. Low-on-resistance (22 mΩ) metal oxide silicon field-effect transistors (CSD17585F5, Texas Instruments) and a low-dropout (120 mV), low-quiescent-current (12 μA) linear regulator (LP5907MFX-3.3, Texas Instruments) with a fixed output voltage of 3.3 V were used to drive LEDs. Passive components with 0201 and 0402 packages further minimized the layout.

### Characterization of the wearable optical device

Current-voltage characteristics were measured using a source meter (Keithley 2400, Tektronix). The electroluminescent spectra were obtained using a spectrometer (USB2000+, Ocean Optics). Thermal measurements were performed using a thermal imaging camera (226s, FOTRIC) and a close-up lens, with a background temperature of 37 °C.

### Animal experiments

Seven-week-old rats were given a high-fat diet for 2 months to produce insulin resistance. Oral glucose tolerance (OGTT) was then tested. The model was considered successful if the blood glucose did not drop below 6 mmol/ml within 2 h. Achieving this setpoint, each rat was injected with 1% stz-1 solution in the tail vein at a dose of 28.5 mg/kg. This procedure produced the rat model of T2D. All rats were fed normal food during the experiment. For DIO mice experiment, after purchasing 16-week-old DIO mice and their control group (fed with normal diet), they were fed with high-fat diet or normal diet for two weeks to adapt to the environment, and then carried out experiments.

### Synthesis of up-conversion microrods

Five microliters of NaOH (8 M) were mixed with 25 ml EtOH and equal oleic acid under continuous agitation for 10 mins at 37°C. A rare-earth solution (consisting of 1 mM, Y3+: Yb3+: Tm3+=75: 25: 0.5) was combined with 4 mL NH4F (2 M) and stirred for 30 mins at 400 rpm to form a homogeneous solution. The mixture was then transferred to a reaction kettle and heated to 180°C for 12 h. The reaction solution was collected and washed by adding the same volume of anhydrous ethanol. After washing, the reagent mixture was centrifuged at 5000 rpm for 3 mins. The precipitate was collected, added to anhydrous ethanol, and resuspended by ultrasound. The microrods were stored in ethanol or PBS.

### Preparation of up-conversion microcapsules (UCMCs)

UCMCs were synthesized using a microfluidic system controlled by two pressure-driven flows. The external liquid was dimethyl silicone oil and the internal liquid contains Poly(ethylene glycol) diacrylate PEGDA-700: PEG-200: 2-hydroxy-2-methyl-1-phenyl-1-propanone (HMPP) : Tris-EDTA (TE) buffer= 2: 4: 1: 3. The up-conversion microrods were dissolved in TE buffer at a final concentration of 10 mg/ml. Internal and external flow rates were controlled by the syringe pump and formed microdroplets in the mixing process. The droplets then crosslinked under the action of ultraviolet light to form stable UCMCs. UCMCs were collected and washed with PBS for use in the in vivo experiments.

### Ethics

All experiments involving animals were performed according to the protocol approved by the Institute of Radiation Medicine Chinese Academy of Medical Sciences and in direct accordance with the Ministry of Science and Technology of the People’s Republic of China on Animal Care Guidelines (GB14925-2010).

## Acknowledgments

This work was sponsored by National Key Research and Development Program of China (2019YFA0906500 and 2017YFA0205104), National Natural Science Foundation of China (31971300, 817719709, 51873150), Tianjin Natural Science Foundation (19JCYBJC28800) and the Key project of Tianjin Foundational Research (JingJinJi) Program, China (19JCZDJC64100).

## Author contributions

X.Z., N.M., H.W. and X.H. contributed to the conception of the study; X.Z., N.M., G.P., T.S., J.L., C.H., H.P. and M.C. performed strain construction, up-conversion synthesis and functional verification experiment; X.Z., N.M. and G.P. performed animal experiment; W.L. and X.H. designed the optogenetic device; X.Z., N.M., H.P. and W.L. analyzed data; J.C., H.W. and X.H. carried out analyses with constructive discussions; X.Z., N.M. and W.L. wrote the manuscript.

X.Z., N.M. and W.L. contributed equally to this work.

## Supporting Information

### Optical modeling of the flexible optogenetic device

A Monte Carlo simulation software ValoMC was used for light penetration depth modeling. The mean free path *l* can be represented by

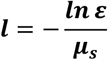

where *ε* is a random variable generated using the Mersenne-Twister algorithm, *μ_s_* is the scattering coefficient of the medium.

The movement of the photon can be represented by

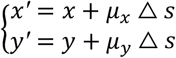

where *μ_x_* and *μ_y_* are the direction cosine values specified by taking the cosine of the angle of the photon with each axis, *Δs* is the distance propagated during the current step. The absorption during photon propagation follows the microscopic Beer-Lambert law, thus the new weight of the photon can be given by

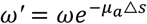

where ω is the initial photon weight, μa is the absorption coefficient of the medium.

The scattering direction can be estimated by the Henyey-Greenstein phase function,

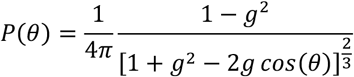

where *g* is the scattering anisotropy factor, so the deflection angle is calculated to be

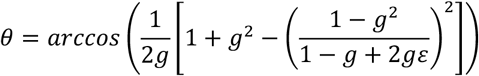

the fluence *Φ_i_* at element *i* can be represented to be

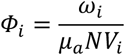

where *ω_i_* is the weight absorbed into the element *i*, *N* is the number of photon packets and *V_i_* is the area of the element.

Briefly, a length of 1 mm on the left boundary was set as the light source, with a cosine launching property. The photon count was set to be 10^6^, and a square area of 1 × 5 mm^2^ was set as the medium to mimic the skin, with an axial resolution of 10 μm. For 460 nm blue light, the absorption coefficient μ_a_ = 0.18 mm^−1^, the scattering coefficient μ_s_ = 42 mm^−1^, the scattering anisotropy factor g = 0.9, and the refractive index n = 1.42. The optical power was set to be 7.25 mW.

**Supplementary Figure 1.**
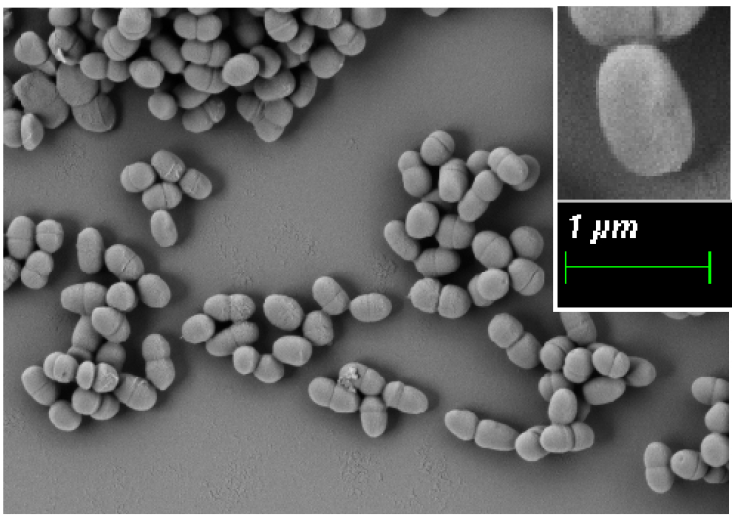
Scanning electron microscope image of engineered *L.lactis*.

**Supplementary Figure 2.**
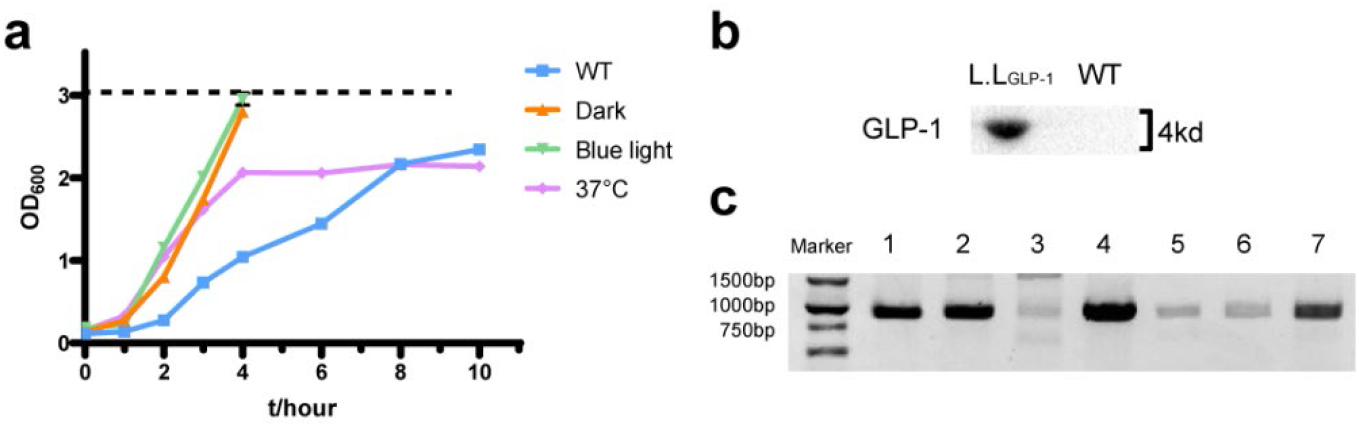
**a**. The growth curve of *L.L_GLP-1_* under dark, blue light and 37°C. **b**. Western blot of GLP-1 secreted by *L.L_GLP-1_* after blue light stimulation. **c**. Colony PCR of *L. lactis* in the feces up to 7 days after gavage.

**Supplementary Figure 3.**
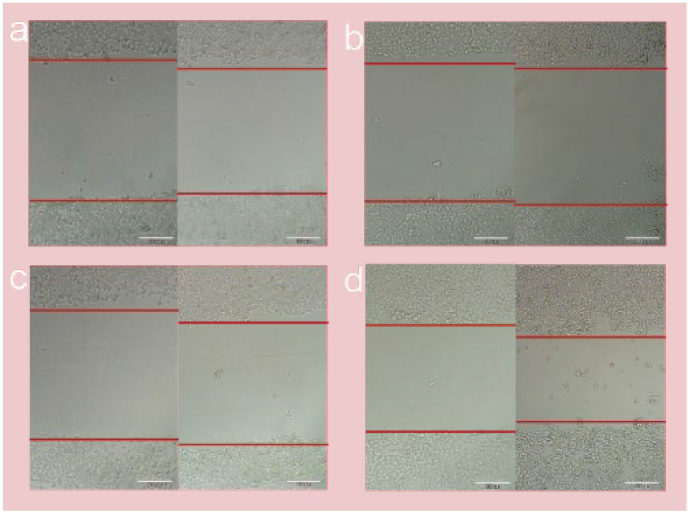
Cell scratch experiment shows GLP-1 promotes proliferation of ins-1 within 24 hours.

**Supplementary Figure 4.**
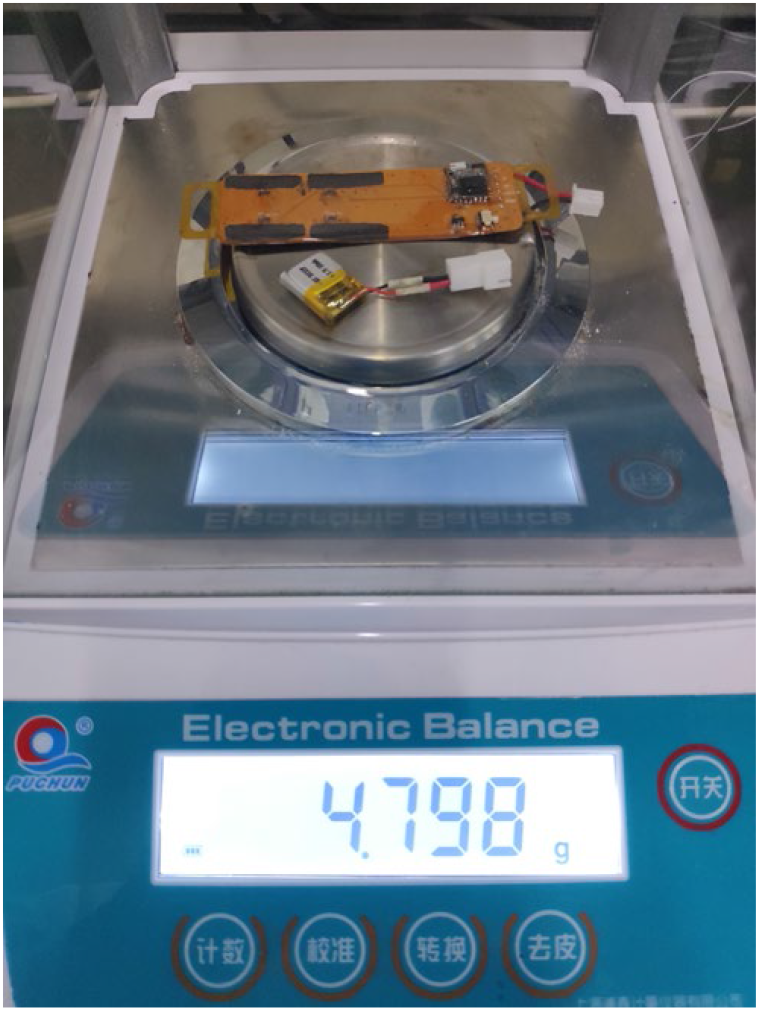
The weight of the battery attached to the LEDs is 4.798 g.

**Supplementary Figure 5.**
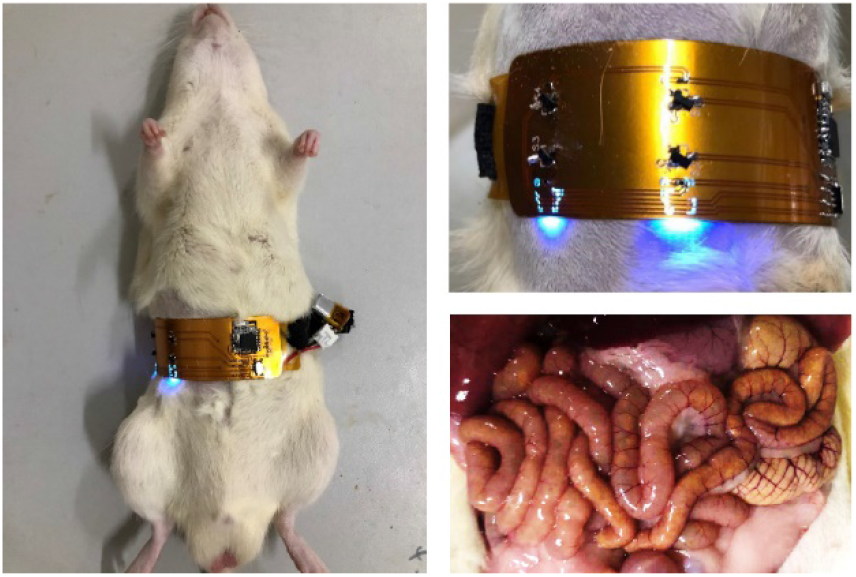
The LED device mainly irradiates the small intestine of rats with its GLP-1 higher absorption capacity.

**Supplementary Figure 6.**
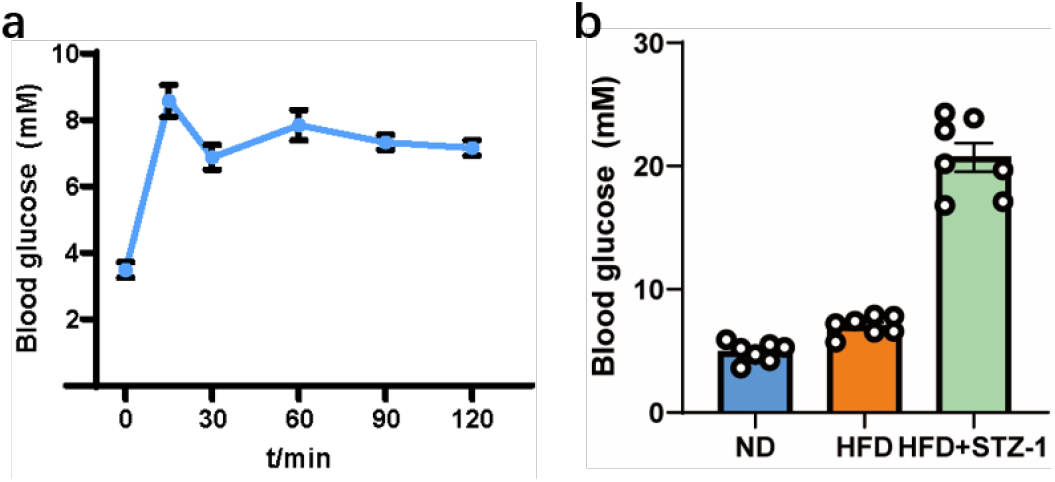
**a**. Before STZ-1 injected, the HFD rats developed insulin resistance after two months of high-fat feeding. **b**. After STZ-1 injected, the concentration of blood glucose of the rats. Sample size: n≥ 5.

**Supplementary Figure 7.**
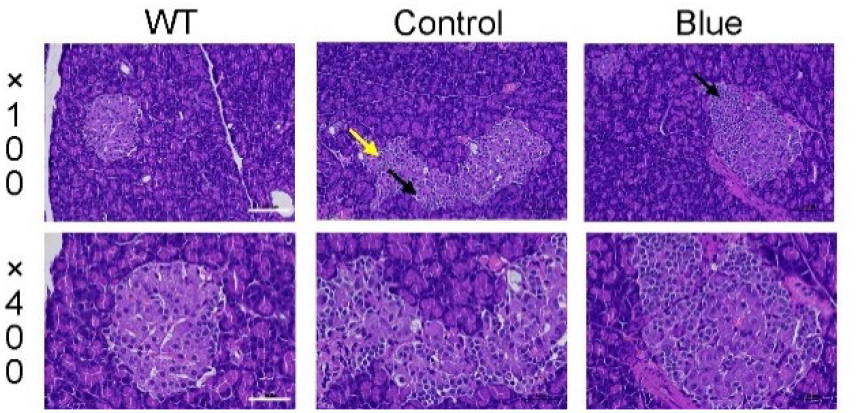
H&E-stained slices of pancreas tissues collected from different groups after treatment. Scale bar: 100 μm (×100); 50 μm (×400).

**Supplementary Figure 8.**
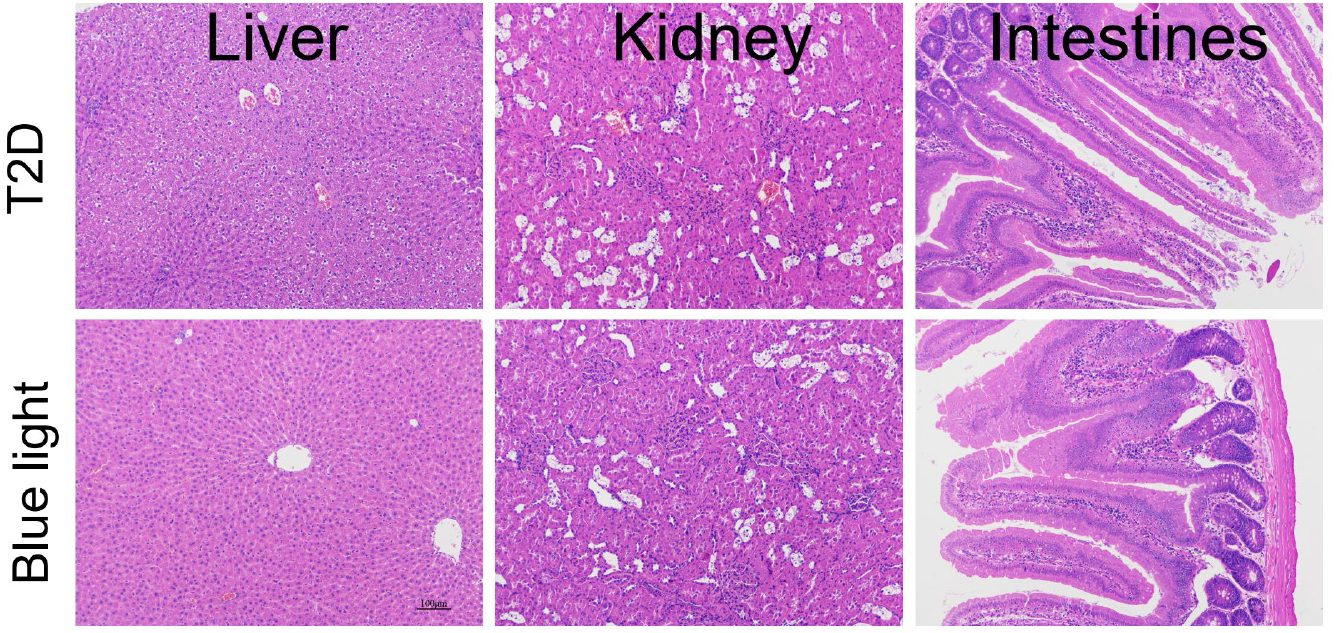
Hematoxylin and eosin (H&E)-stained slices of liver, kidney and intestines of rats in the control group and blue light group. No significant tissue damage was found.

**Supplementary Figure 9.**
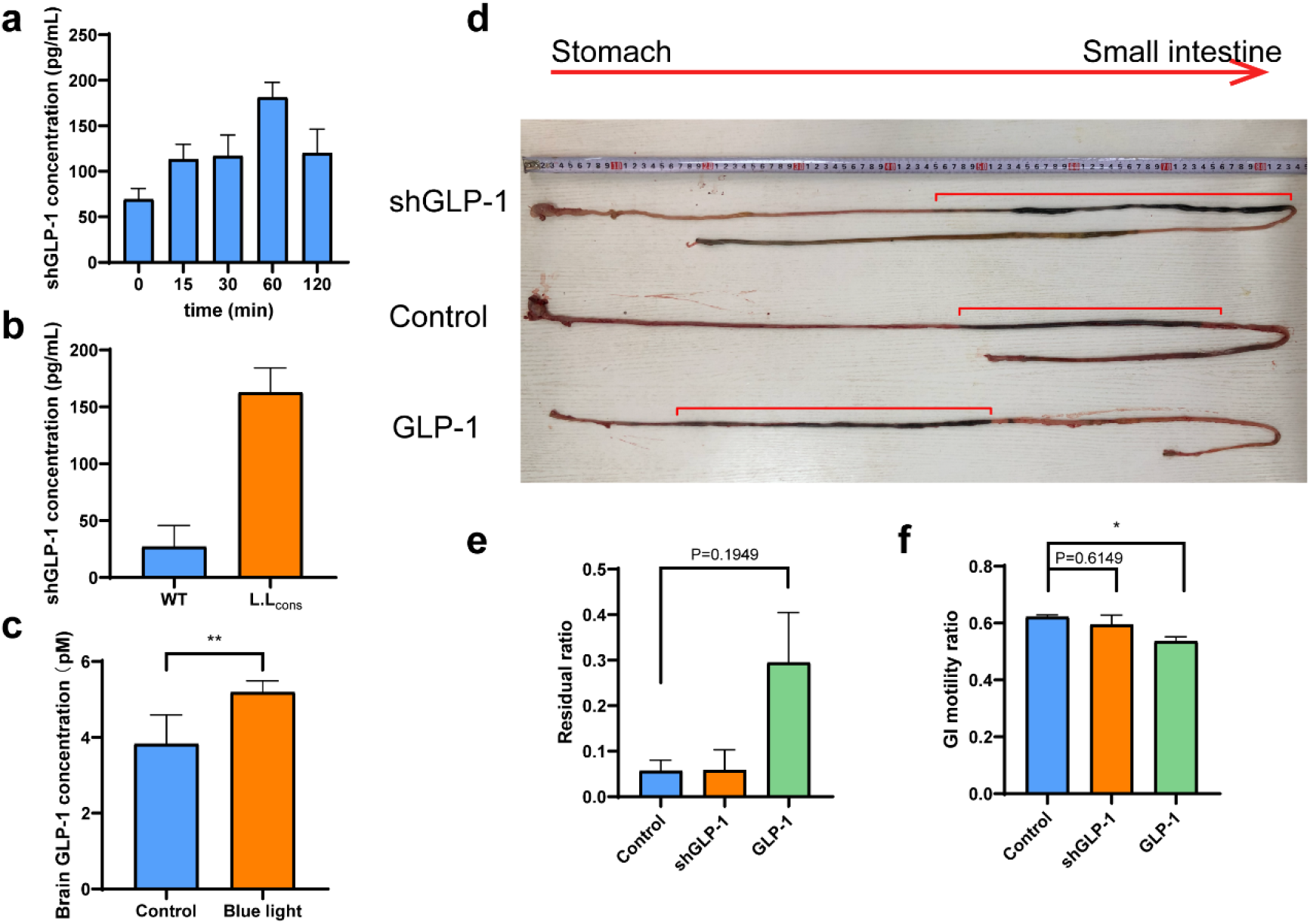
**a**. and **b**. shGLP-1 secreted by *L.L_sh_* under blue light and *L.L_cons_*. **c**. GLP-1 detected in the brain of rats under blue light stimulation. **d**. GI motility and gastric emptying. (n=3) The red line showed mash mixed with carbon powder in the intestine. **e**. Proportion of residual mash in the stomach. **f**. Proportion of the distance the mash travels in the gut.

**Supplementary Figure 10.**
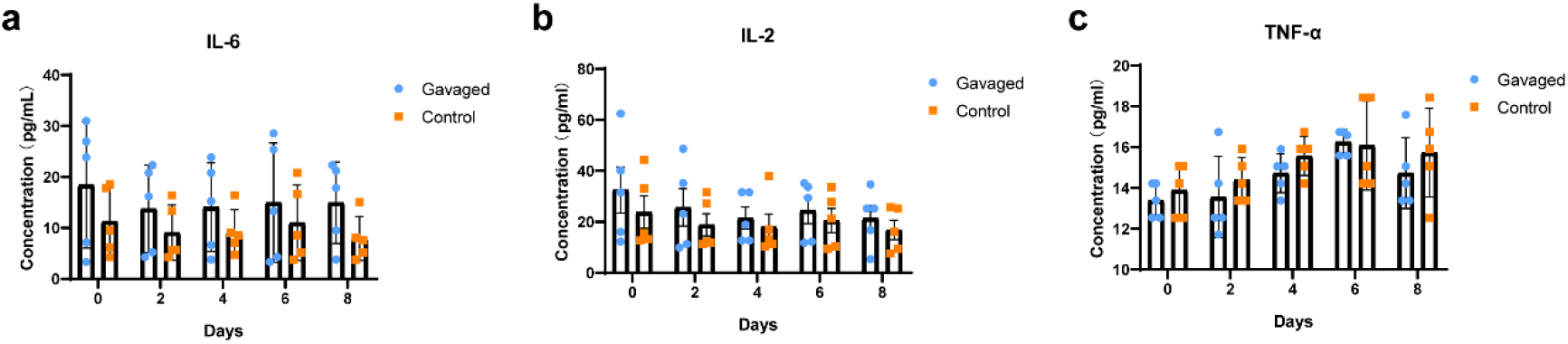
Immune factors IL-6 **(a)**, IL2 **(b)** and TNF-α **(c)** in serum tested by ELISA after *L.latics* gavage.

**Supplementary Figure 11.**
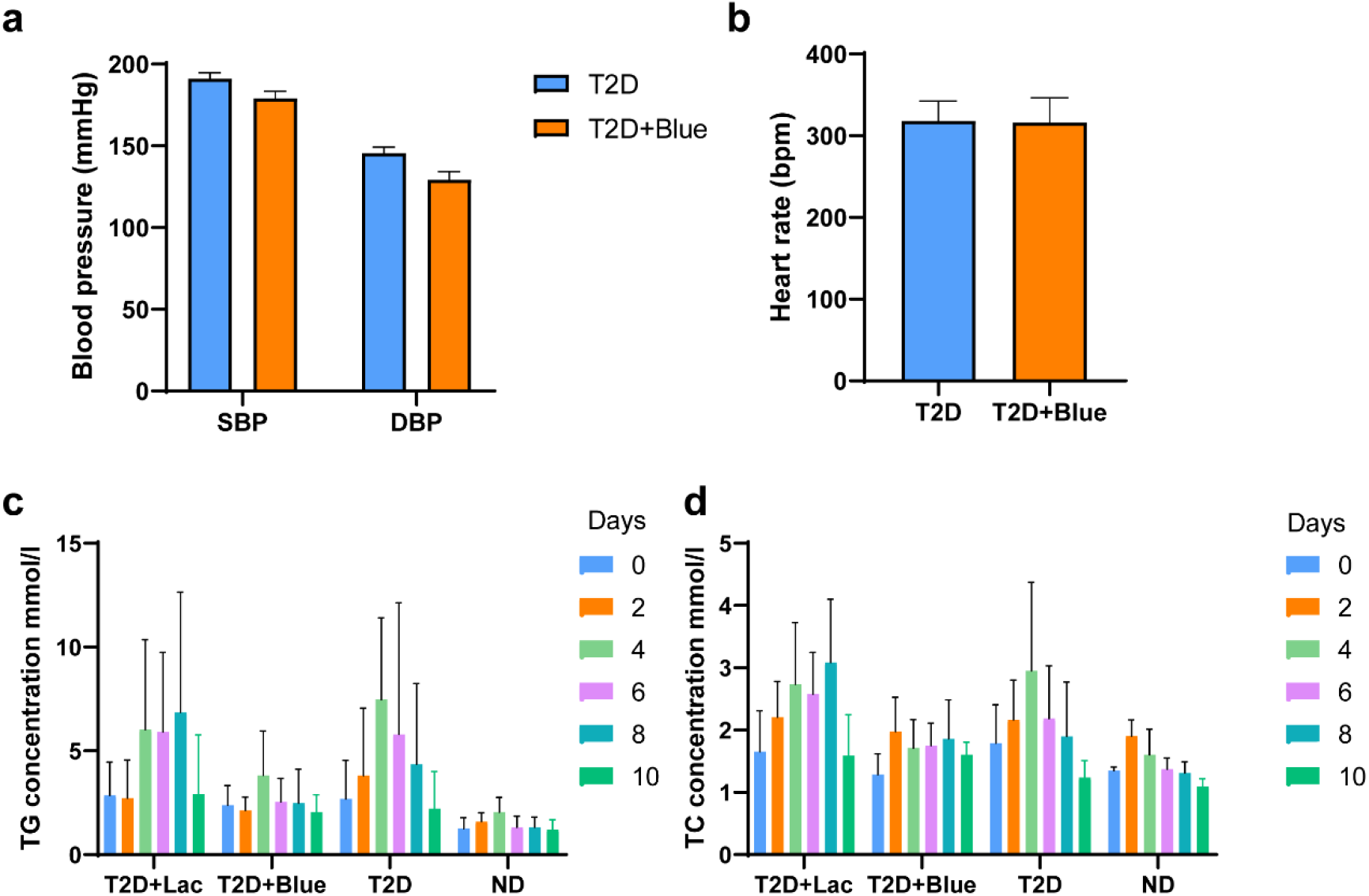
Physiological effect of shGLP-1 secretion. **a**. Blood pressure change of T2D rats after 30 days’ treatment. **b**. Heart rate change of T2D rats after 30 days’ treatment. **c** and **d**. TG and TC concentration tested by ELISA in the first ten days of treatment. (n≥5).

**Supplementary Figure 12.**
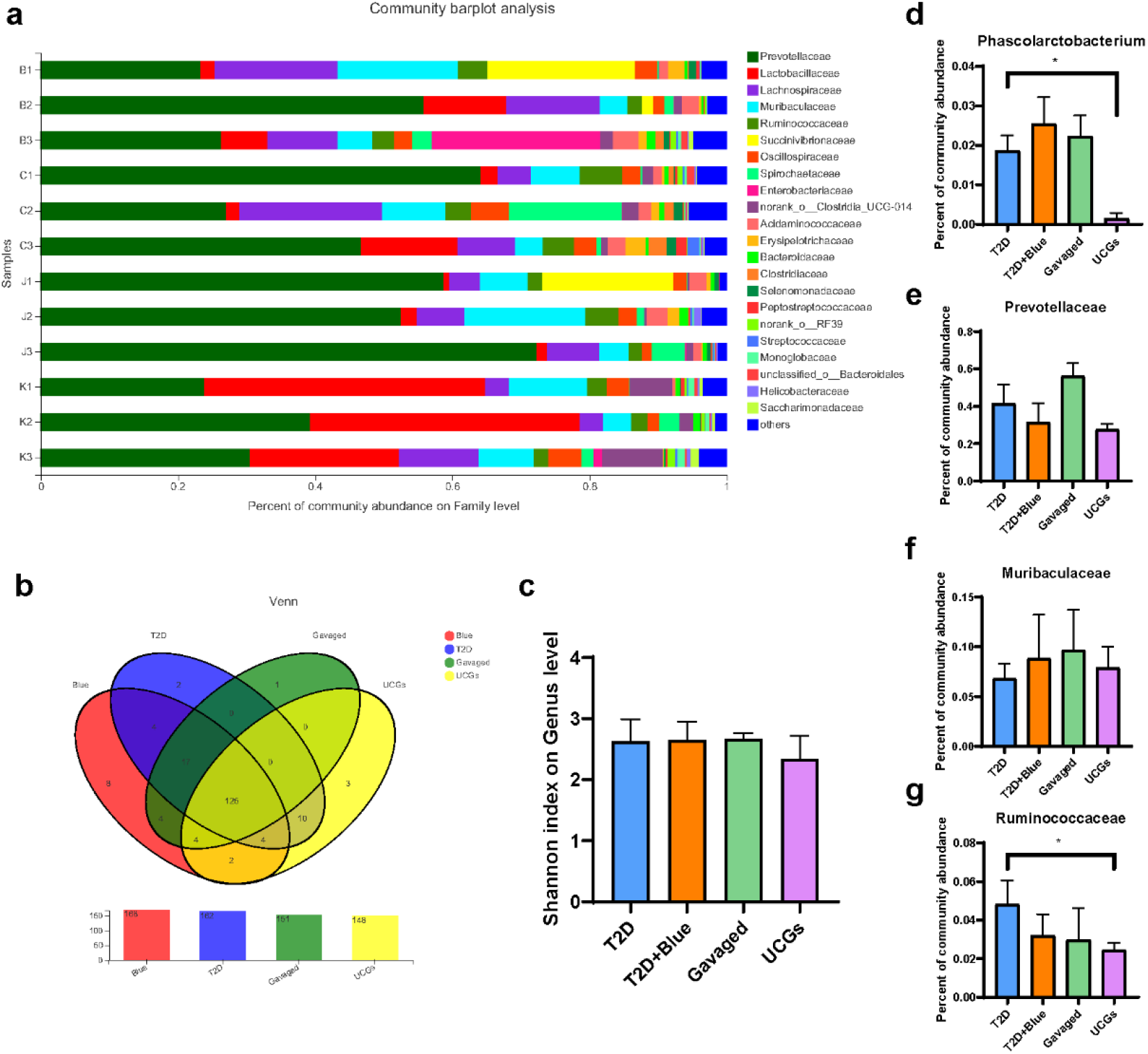
Effect of the OBFE system on rats. **a**. Percent of community abundance of rats treated with blue light (B1-B3), control (C1-C3), *L.latics* (J1-J3) or UCGs (K1-K3) on Family level. **b**. Venn diagram analysis of intestinal flora. **c**. Shannon index reflecting community diversity, there were no significant differences between the groups. **d-e**. Percent of community abundance of Prevotellaceae, Muribaculaceae, Phascolarctobacterium and Ruminococcaceae.

**Supplementary Figure 13.**
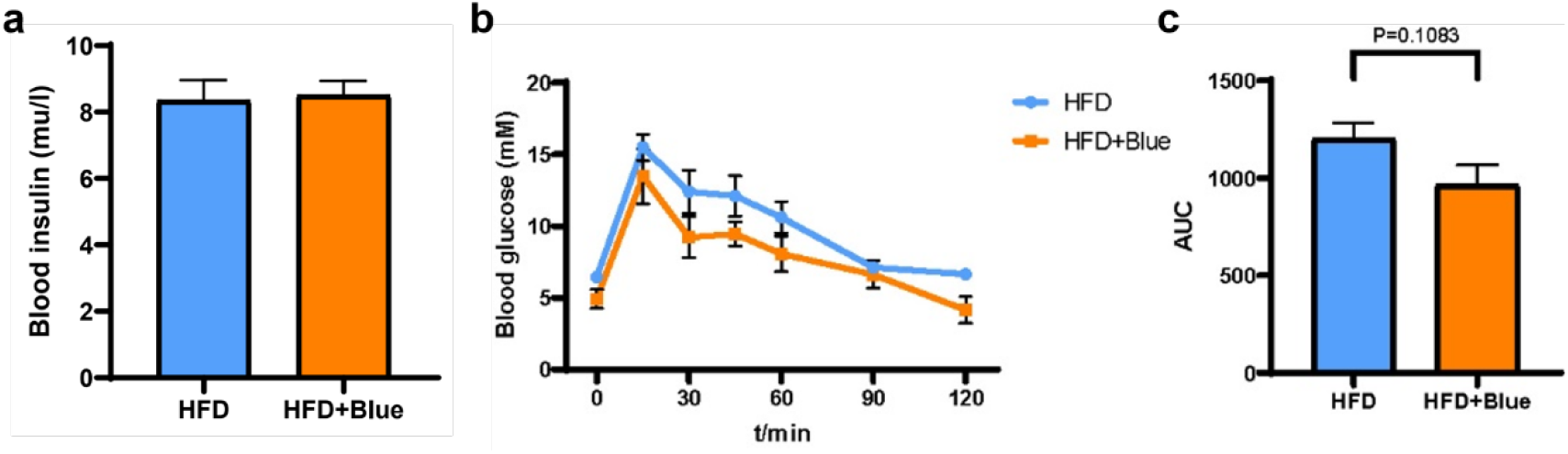
**a**. Insulin concentration change during blue light stimulation in DIO mice. **b**. Glucose tolerance of DIO mice on the 3^rd^ week of treatment. **c**. The area under the curve in **b**. Sample size: n≥ 5. Data are mean ± SEM. P values were calculated by Student’s t test.

**Supplementary Figure 14.**
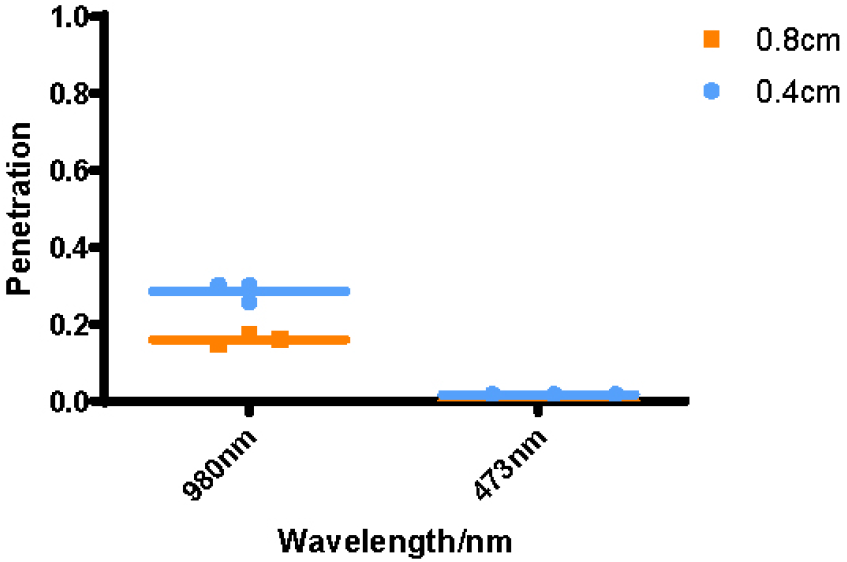
Biological tissue penetration contrast between blue light and NIR.

**Supplementary Figure 15.**
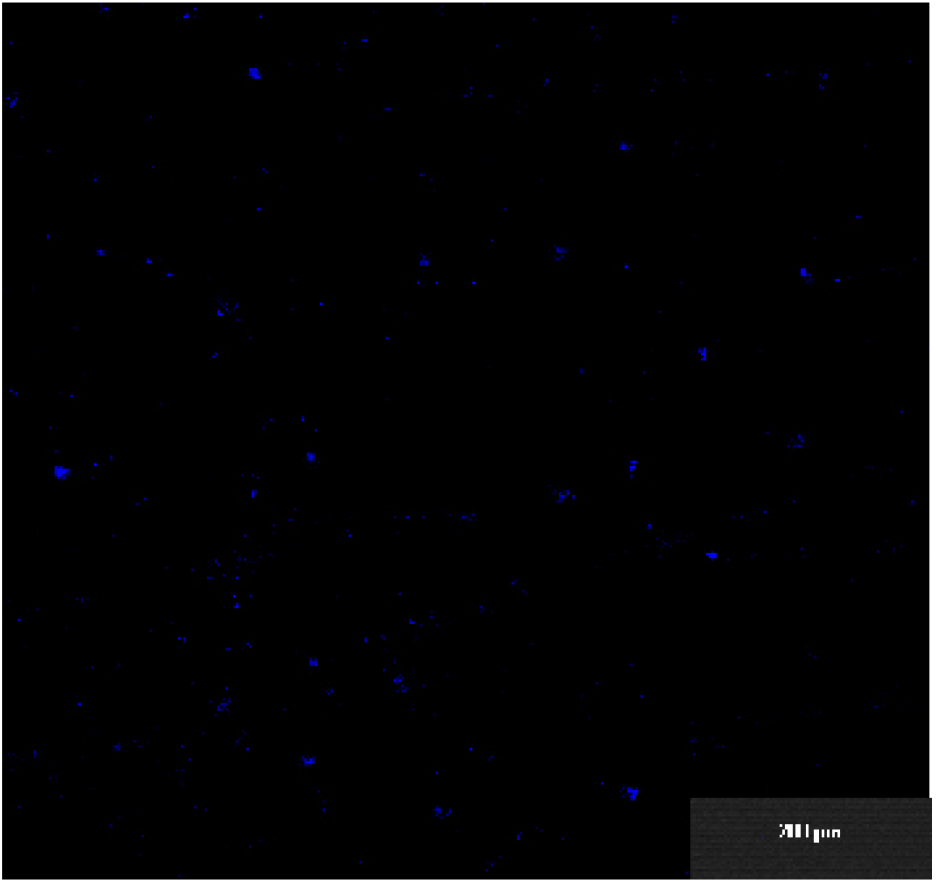
Confocal microscope fluorescence photograph of up-conversion microrods under 980 nm light excitation.

**Supplementary Figure 16.**
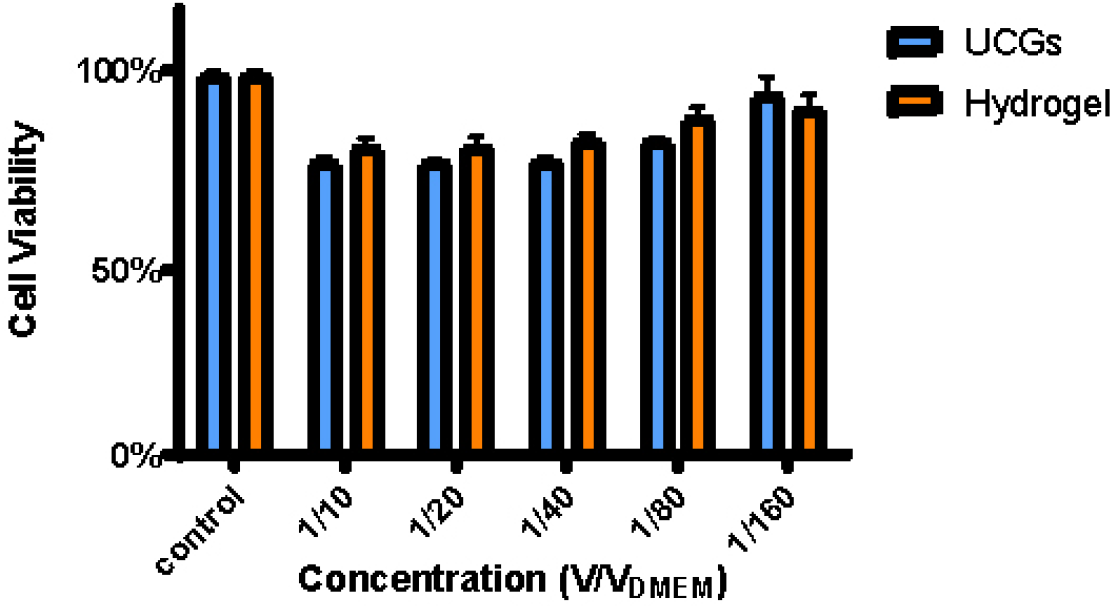
Biosafety experiment (CCK-8) of UCGs using hela cell.

**Supplementary Figure 17.**
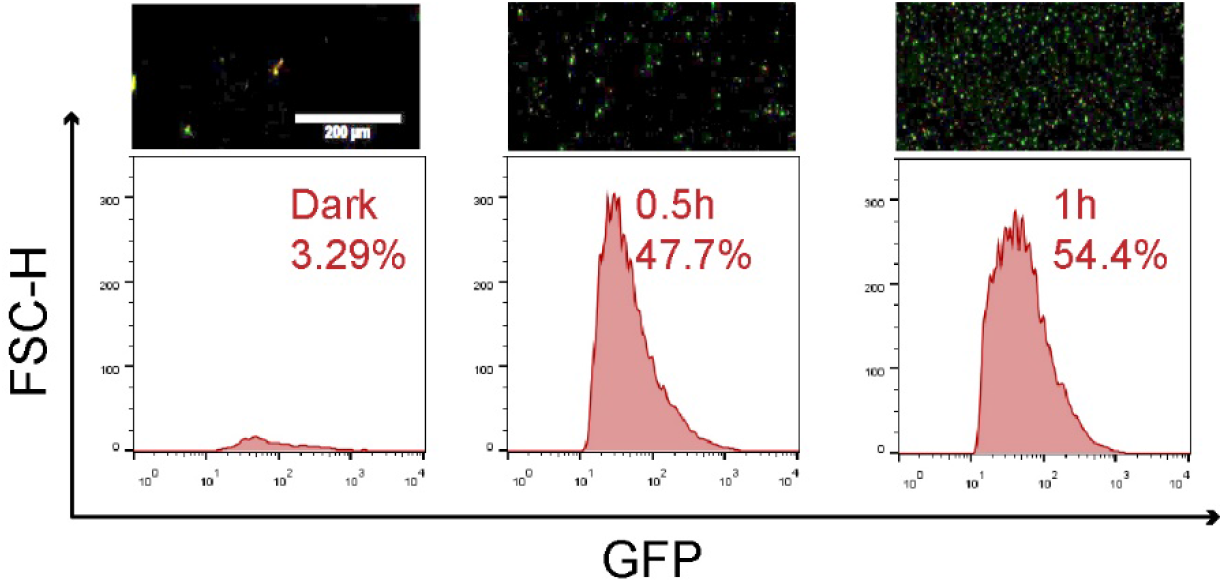
Microscope fluorescence images and flow cytometric analysis of *L.L_GFP_* colonized in the intestinal tract induced by NIR.

